## Supplementary figures and images for "*ZNF423* patient variants, truncations, and in-frame deletions in mice define an allele-dependent range of midline brain abnormalities"

### S1_Fig

S1_Fig. Related to Figure 2

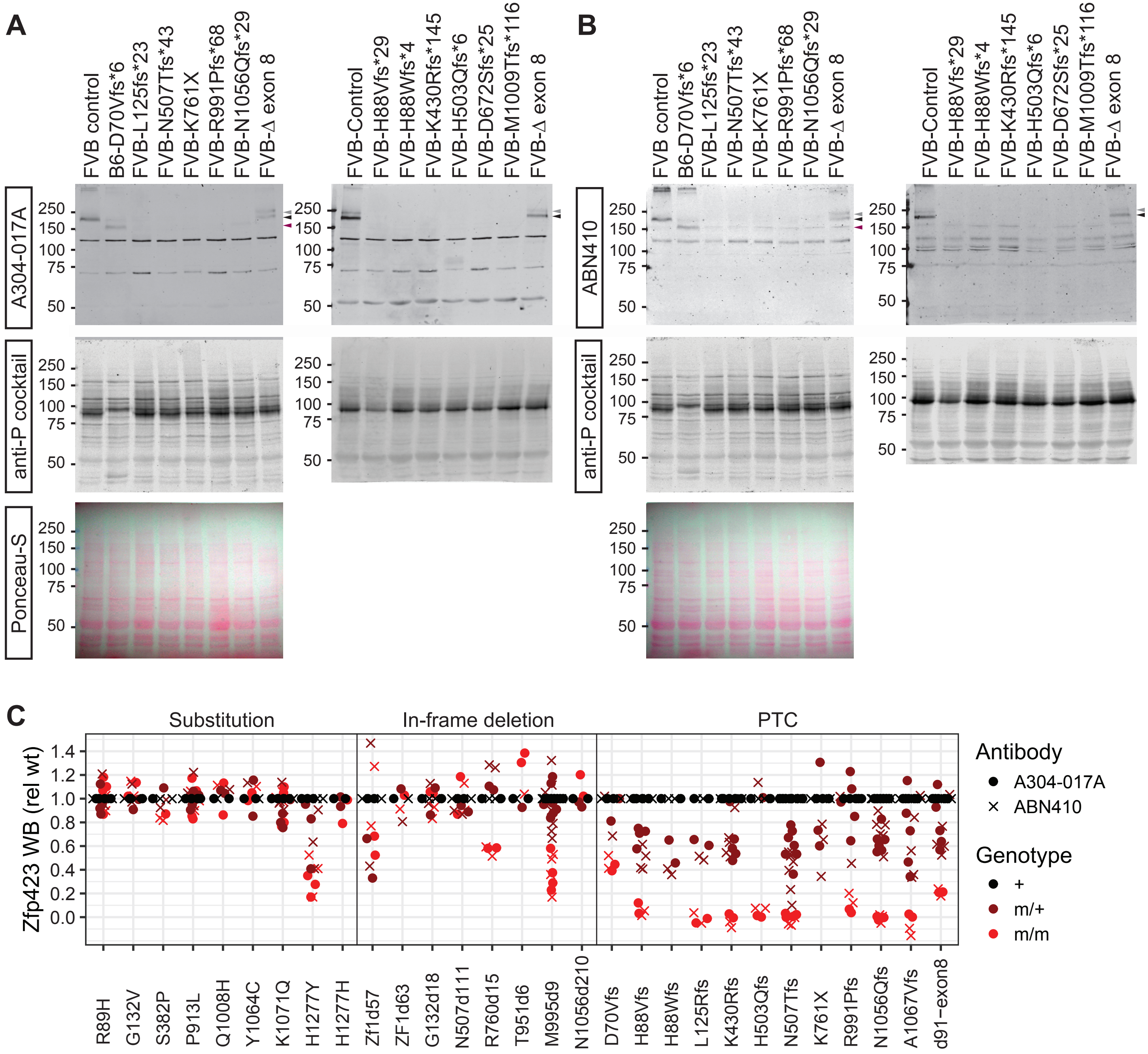

### S3_Fig

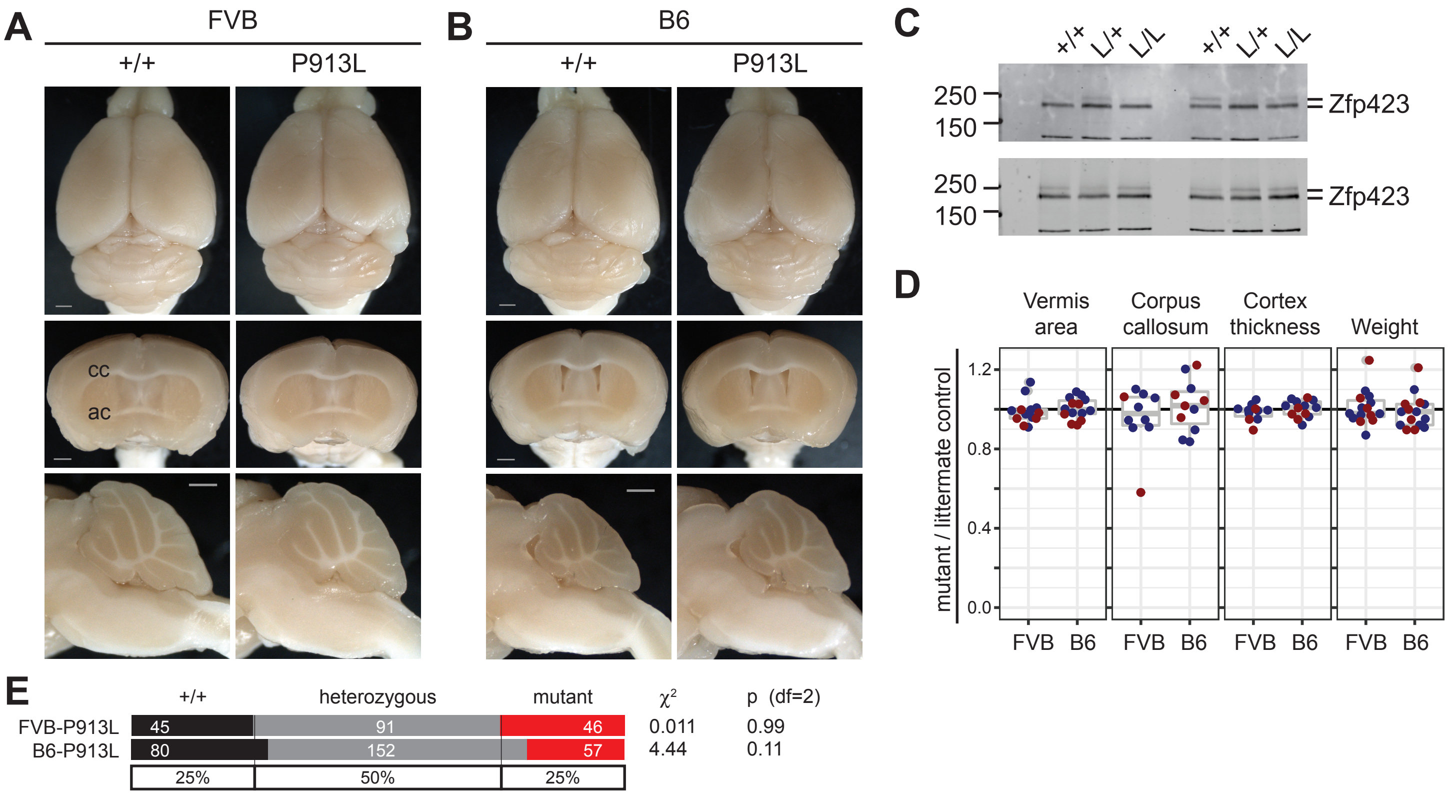

### S4_Fig

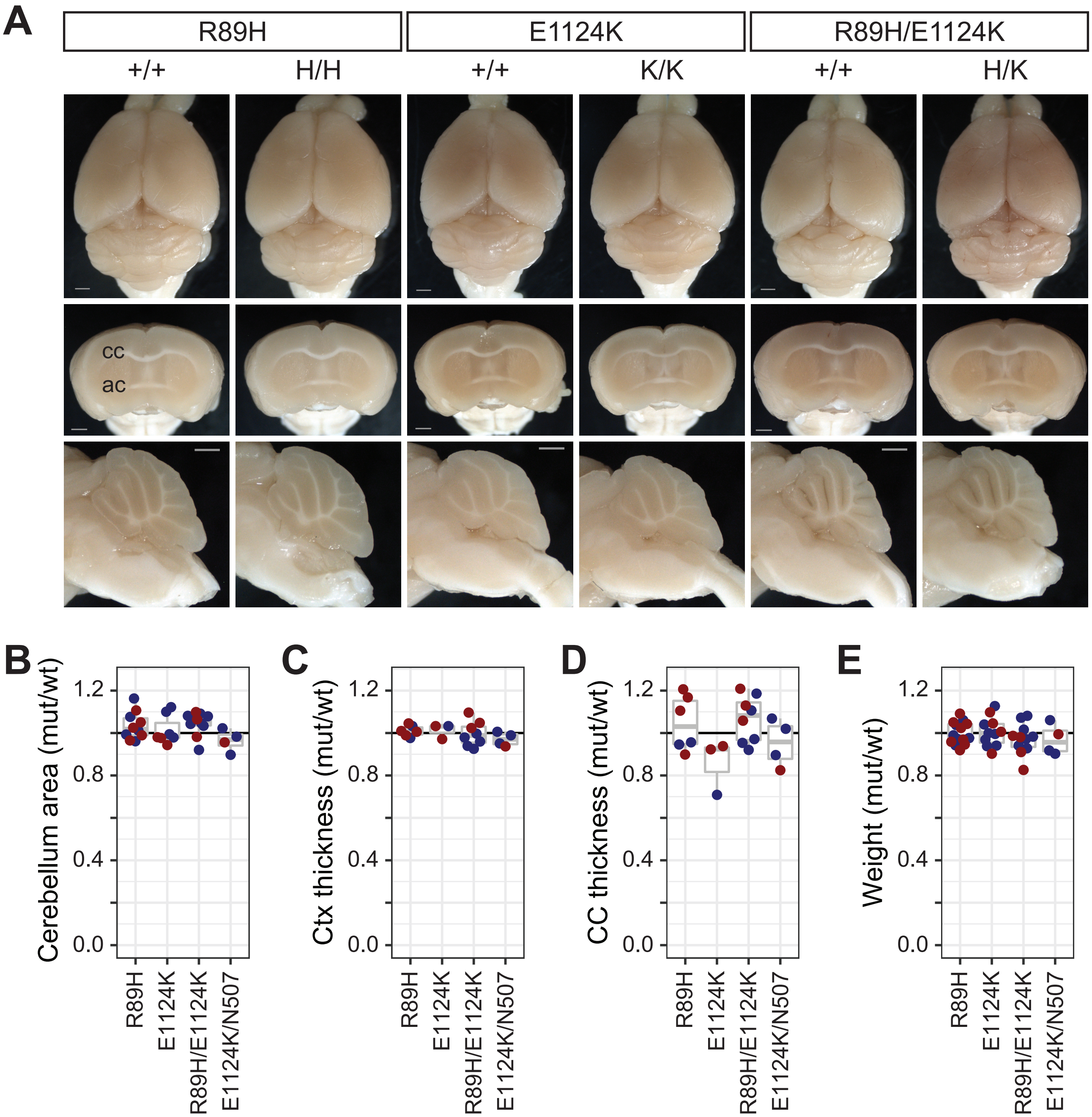

### S5_Fig

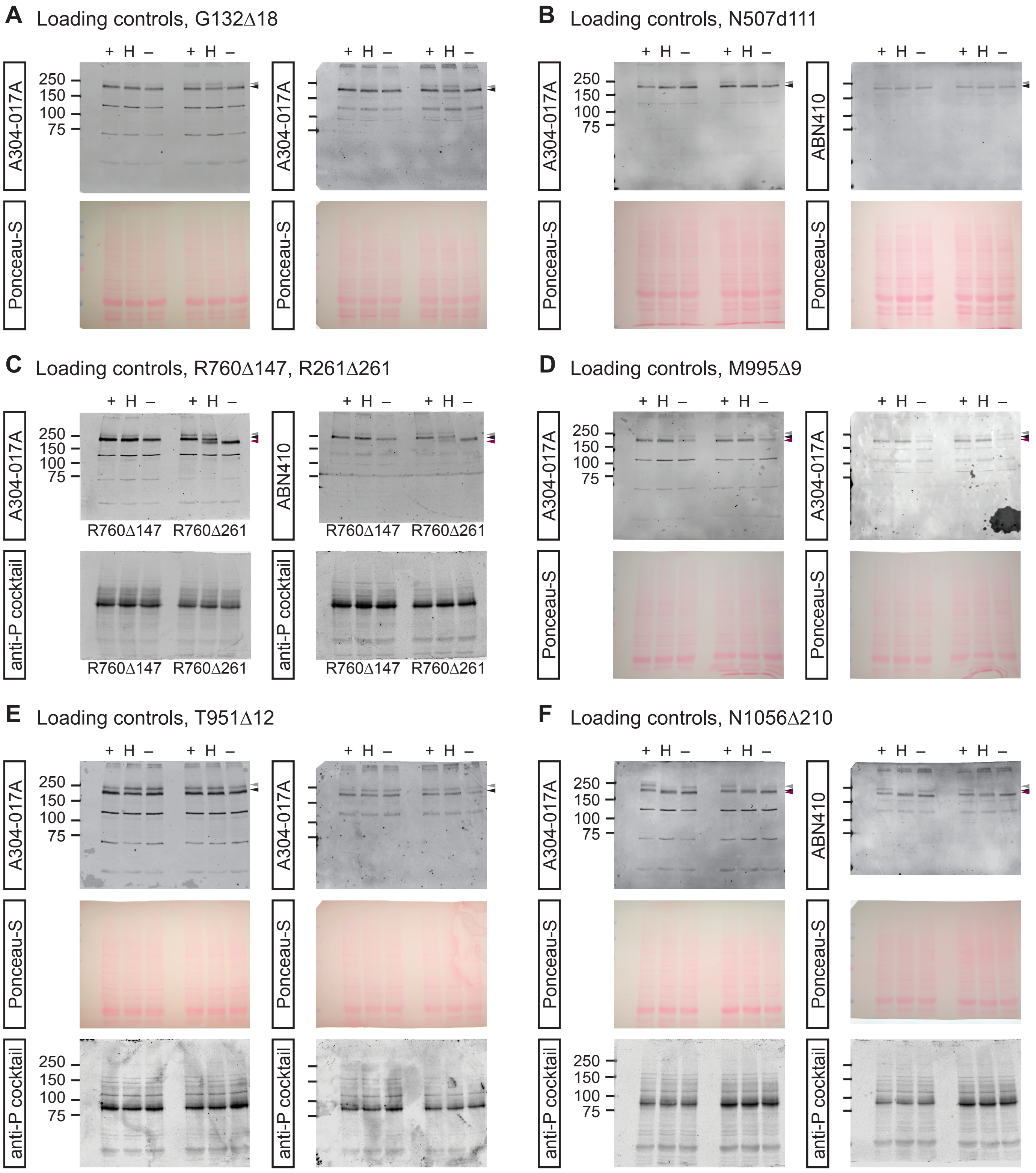
