## Supplementary material for "*ZNF423* patient variants, truncations, and in-frame deletions in mice define an allele-dependent range of midline brain abnormalities": S2_Fig

**A** Loading controls, Figure 3C

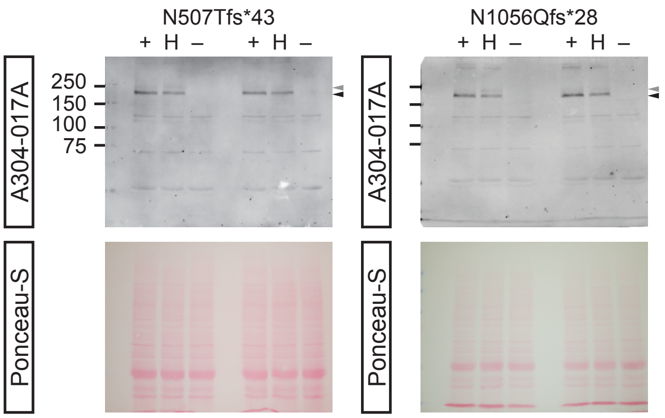


**B** Loading controls, Figure 4A

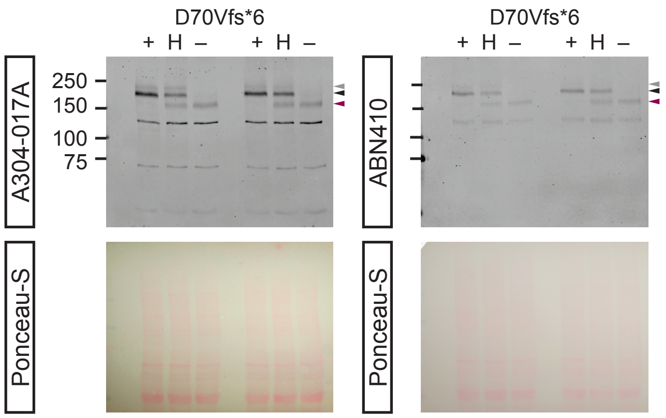


**C** Loading controls, Figure 4B

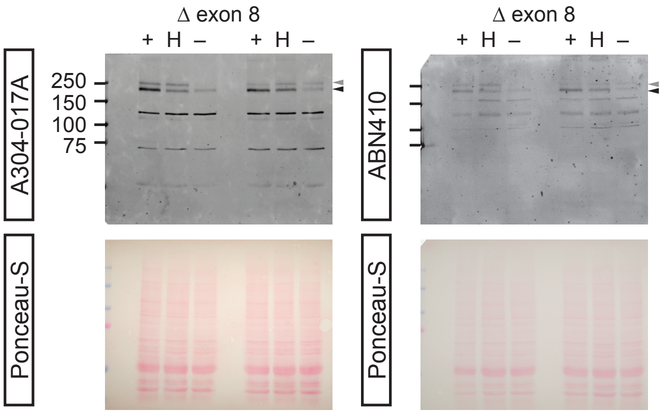


**D** Loading controls, Figure 5C

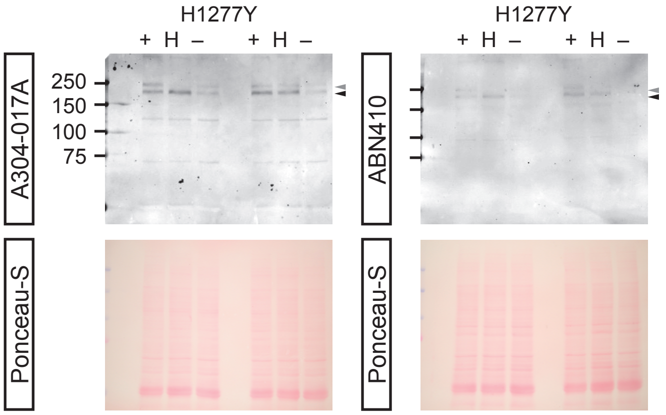


**E** Loading controls, Figure 5S1C

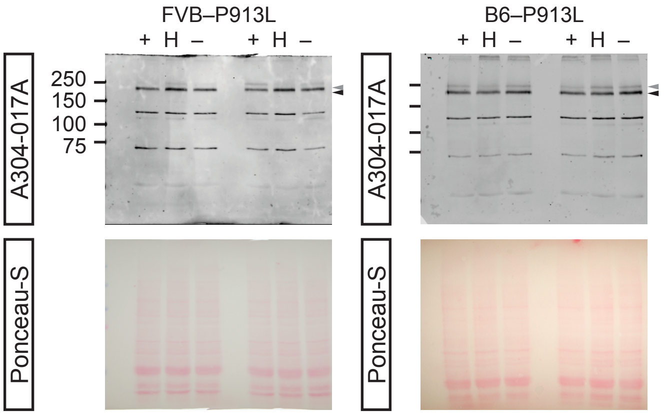
